# Scalable Culture of Mammalian Cells in RGD Peptide-Modified Alginate Hydrogel Microtubes

**DOI:** 10.1101/2025.04.22.650014

**Authors:** Ou Wang, Li Han, Cheng Dong, Liangqi Xie, Aijun Wang, Shue Wang, Yuguo Lei

**Affiliations:** Department of Chemical and Biomolecular Engineering, University of Nebraska, Lincoln, Nebraska, USA; Department of Biomedical Engineering, Pennsylvania State University, PA, USA; Cancer Biology and Infection Biology, Lerner Research Institute, Cleveland Clinic, Cleveland, OH, USA; Department of Biomedical Engineering, University of California, Davis, Davis, CA, USA; Center for Surgical Bioengineering, Department of Surgery, University of California, Davis, School of Medicine, Sacramento, CA, USA; Department of Chemistry, Chemical and Biomedical Engineering, University of New Haven, CT, USA; Huck Institutes of Life Sciences, Pennsylvania State University, PA, USA

**Keywords:** Cell cultured meat, biomanufacturing, bioreactor, alginate hydrogel tube

## Abstract

Traditional livestock farming is resource-intensive and environmentally unsustainable, necessitating alternative methods for meat production. Cell cultured meat, produced by expanding and differentiating animal cells, offers great potential for substituting for conventional animal meat. Nevertheless, it is still limited by the scalability and efficiency of current cell culture technologies. In this study, we developed an RGD peptide-modified alginate hydrogel microtube microbioreactor (AlgTubes) to support the scalable culture of anchor-dependent cells, such as myoblasts and adipocytes, for cell-cultured meat production. AlgTubes provide a cell-friendly 3D microenvironment that enhances cell viability, growth, and yield while overcoming limitations of conventional bioreactors, such as shear stress, aggregation, and diffusion constraints. We successfully expanded mouse (C2C12) and quail (QM7) myoblasts in AlgTubes, achieving cell densities exceeding 1.0 × 10⁸ cells/mL, far surpassing traditional stirred-tank bioreactors. Differentiation resulted in the formation of mature myotubes. Co-culturing myoblasts with mesenchymal stem cells or fibroblasts further improved yield and viability, particularly under differentiation conditions. By significantly increasing cell culture density, AlgTubes can substantially reduce culture volume, lowering labor requirements, reagent costs, equipment needs, facility space, and manufacturing expenses.

## Introduction

Based on the Food and Agriculture Organization (FAO) data, the global population is expected to reach 9 billion by 2050. Accordingly, 70% more food will be needed to feed this population. Meat is a significant part of human food. Currently, meat is obtained from natural animals or livestock raised through traditional farms or modern factories^1–3^. Livestock farming requires large amounts of water and land and will eventually reach a limitation. Additionally, farming causes significant environmental and animal welfare issues^4,5^. Alternative approaches that can produce meat efficiently and environmentally friendly are highly needed.

Cultured meat, produced by culturing animal cells to replicate conventional meat’s organoleptic and nutritional properties, holds significant promise as a sustainable meat substitute^6–8^. In this process, animal cells are expanded to large numbers and differentiated into myocytes, which are assembled into muscle tissue. However, this approach remains at the proof-of-concept stage, with several key technological challenges must be addressed to enable further development. One of the primary challenges is achieving mass cell production, which requires overcoming limitations in the efficiency, cost, and scalability of existing cell culture technologies. It is estimated that producing 1 kg of muscle tissue requires approximately 3 × 10¹² cells. Current culture methods, including two-dimensional (2D) flasks and three-dimensional (3D) stirred tank bioreactors (STRs), are incapable of reaching this scale. For example, STRs typically yield only ∼2 × 10⁶ stem cells per milliliter, with the resulting cell mass occupying merely ∼0.4% of the culture volume^9–11^. Alternative methods, such as culturing cells within 3D hydrogel scaffolds, also result in low yields. Consequently, the relatively low working cell densities (10⁵–10⁶ cells/mL) and limited working volumes (∼50 L) of current bioreactor systems present significant barriers to scalability.

Previously, our lab developed a new cell culture system termed AlgTubes^20–32^. This method cultivates cells in hollow, microscale alginate hydrogel tubes. AlgTubes provide a cell-friendly microenvironment, resulting in paradigm-shifting improvements in cell viability, growth rate, yield, culture consistency, and scalability. When culturing human stem cells, we have achieved up to 4000-fold expansion per passage and 5×10^8^ cells/mL volumetric yield, which is ∼250 times the current state-of-the-art.

Their high growth rate and cell density profoundly impact large-scale cell production. For instance, producing 10¹² human pluripotent stem cells-derived cardiomyocytes would require ∼1,000 liters of culture volume using stirred-tank bioreactors. In contrast, the same production can be achieved with just 2 liters of AlgTubes. This dramatic reduction in culture volume significantly reduces labor, reagent costs, equipment needs, facility space, and overall manufacturing expenses, making large-scale production feasible and practical. These results highlight the transformative potential of hydrogel tube microbioreactors for stem cell culture. However, AlgTubes do not support the growth of anchor-dependent stem cells, as they lack adhesion points. In this work, we re-engineered the AlgTubes to support anchor-dependent cells and studied it for culturing animal myoblasts to produce meat.

## Materials and Methods

### Cell Lines and Materials

Mouse bone marrow mesenchymal stem cell line (D1, CRL-12424), mouse myoblast cell line (C2C12, CRL-1772), quail myoblast cell line (QM7, CRL-1962), mouse beige adipocyte cell line (X9, CRL-3282), mouse fibroblast cell line (3T3, CRL-1658) were purchased from ATCC and maintained as instructed by ATCC. Briefly, C2C12, D1, and 3T3 were cultured in DMEM and supplemented with 10% FBS. QM7 were cultured in medium 199 supplemented with 10% TPB and 10% FBS. X9 were cultured in DMEM/F12 supplemented with 15% FBS and 2.36 mM L-glutamine. All cell culture medium was refreshed every 2-3 days. To induce C2C12 and QM7 myotube formation, the culture medium was supplemented with 10% horse serum instead of fetal bovine serum.

Additionally, the following chemicals and reagents were also used for this study: sodium alginate (cat # 194-13321, 80∼120cp, Wako Chemicals), sodium hyaluronate: (cat # HA 700K-5, Lifecore Biomedical), tryptose phosphate broth (TPB, cat # 18050039, Life Technologies), Dulbecco’s Modified Eagle’s Medium (DMEM, cat # 30-2002, ATCC), Medium 199 with Earle’s BSS (cat # 12119F, Lonza), DMEM/F12 (cat # 30-2006, ATCC), Fetal Bovine Serum (FBS, cat # 10437028, Gibco), L-Glutamine (cat # 25030081, Gibco), propidium iodide (cat # 195458, MP Biomedicals, LLC), Vybrant multi-color cell-labeling kit (cat # V22889, Molecular Probes), MF20 antibody (cat # MAB4470, R&D systems), RGD peptide (Genscript), Alginate Lyase (cat # A1603, Sigma).

### Modifying Alginates with RGD peptides

2% alginate was dissolved in 0.1N NaOH and reacted with divinyl sulfone or DVS (1:3 molar ratio between OH group and DVS) for 15 minutes. Dialysis was done to remove excessive DVS. 20% to 30% of the OH groups in alginate polymers were modified with DVS. RGD peptides containing a C-terminal cysteine were reacted with alginate-vs under alkaline conditions to make alginate-RGD. 10% of the modified OH groups were reacted with RGD peptides. Alginate-RGDs were mixed with unmodified alginates to make a 2% alginate solution to process alginate hydrogel tubes.

### Processing alginate hydrogel tubes (AlgTubes)

A custom-made micro-extruder was used to process AlgTubes. A hyaluronic acid (HA) solution containing single cells and an alginate/alginate-RGD solution were pumped into the central and side channels of the micro-extruder, respectively, to form coaxial core-shell flows that were extruded into a CaCl_2_ buffer (100 mM) to form AlgTubes. Subsequently, CaCl_2_ buffer was replaced by the cell culture medium. Detailed methods for processing AlgTubes can be found in our previous publications^20–32^.

### Culturing cells in AlgTubes

For a typical cell culture, 20 µL of cell solution in AlgTubes were suspended in 3 mL medium in a 6-well plate. Cells were seeded at the density of 1-2×10^6^ cells/mL hydrogel tube space. 2% alginate modified with 1 mM RGD peptide was used. The hydrogel tube diameter was 200 – 300 µm with shell thickness around 30-70 µm. To passage cells, the medium was removed, and alginate hydrogels were dissolved with 0.5mM EDTA and 100 µg/ mL alginate lyase for 10 min at 37 °C. Cell mass was collected by centrifuging at 100 g for 5 min and treated with 0.25% trypsin-EDTA at 37 °C for 10 min and dissociated into single cells. Digestion was neutralized by a complete cell culture medium.

### Immunocytochemistry

Cells cultured on 2D were fixed with 4% paraformaldehyde (PFA) at room temperature for 15 min, permeabilized with 0.25% Triton X-100 for 10 min, and blocked with 5% donkey serum for 1 h before incubating with primary antibodies in DPBS + 0.25% Triton X-100 + 5% donkey serum at 4 °C overnight. After washing, secondary antibodies were added and incubated at room temperature for 2 h followed by incubating with 10 mM 4’,6-Diamidino-2-Phenylindole, Dihydrochloride (DAPI) for 10 min. Cells were washed with DPBS three times before imaging with Fluorescent Microscopy. For 3D fibrous cell mass immunostaining, cell mass was fixed with 4% PFA at 4 °C overnight. 40 μm thick tissue sections were obtained via cryosection. The sections were washed with DPBS three times and stained as the 2D cell cultures. Alternatively, the cell mass was directly incubated with the primary antibody at 4 °C for 48 h after fixed and washed. After extensive wash, a secondary antibody was added and incubated at 4 °C for 24 h followed by incubating with 10 mM DAPI for 10 min and imaged with Fluorescent Microscopy.

### Statistical analysis

The data are presented as the mean ± SEM. Data was analyzed using GraphPad Prism 8 statistical software and shown as mean ± standard error of the mean. P value was determined by one-way analysis of variance (ANOVA) for comparison between three or more groups or unpaired two-tailed t-tests for two groups. The significance levels are indicated by p-value, *: p<0.05, **: p<0.01, ***: p<0.001.

## Results

### The RGD-AlgTube culture system

We modified our alginate hydrogel tubes (AlgTubes) with RGD peptides so that anchor-dependent cells could be cultured in AlgTubes. Briefly, the sodium alginate was dissolved in 0.1 N NaOH and reacted with divinyl sulfone to generate VS groups on the alginate polymers^33^. The RGD peptides with free-SH groups then reacted with these VS groups via Michael addition reaction (**Figure 1A**). Alginate-RGD was mixed with unmodified alginate to produce a 2% alginate solution with the final 1 mM RGD concentration, which was used to process AlgTubes. To process AlgTubes, a cell solution and an alginate-RGD solution were pumped into the central and side channels of a custom-made micro-extruder, respectively, to form a coaxial core-shell flow that was extruded through a nozzle and into a CaCl_2_ buffer (100 mM) (**Figure 1B and 1C**). Ca^2+^ ions crosslinked the shell alginate flow in seconds to form an alginate hydrogel tube. Subsequently, cells were grown in the tubes suspended in the cell culture medium.

**Figure 1.**
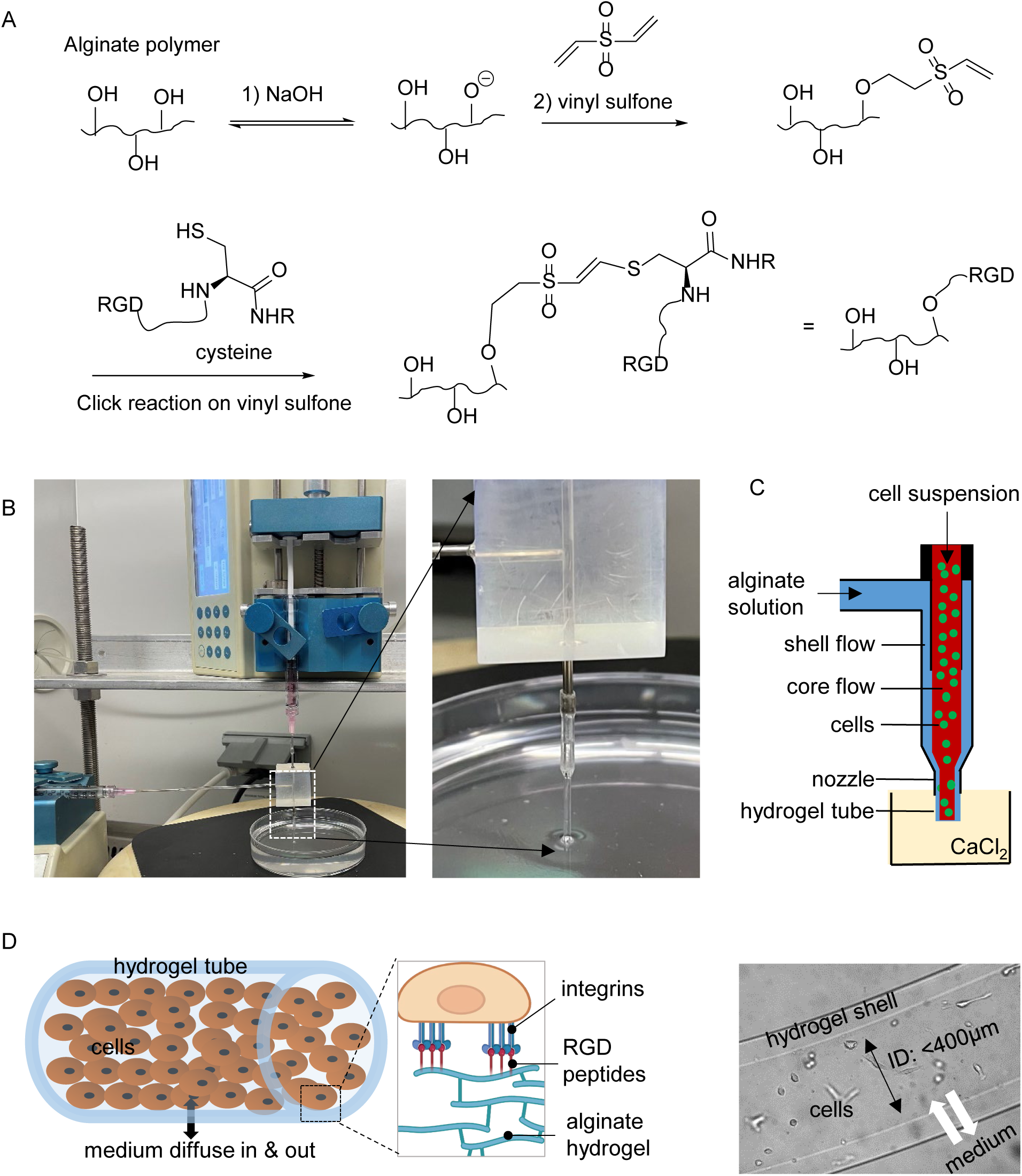
Overview of alginate hydrogel tubes modified with RGD peptides (AlgTubes). (A) Chemistry for making alginate-RGDs. (B, C) Process RGD-AlgTubes. A cell suspension and an alginate solution is pumped into the central channel and side channel of the microextruder to form coaxial core-shell flows that are extruded through the nozzle into a CaCl_2_ buffer. The shell alginate flow is crosslinked by Ca^2+^ ions to form an alginate hydrogel tube instantly. (D) Illustration of cell microenvironment in AlgTubes.

The AlgTubes system is designed to provide cells with a friendly microenvironment (**Figure 1D**). First, RGD peptides allow cells to attach to a substrate and proliferate. Second, oxygen, nutrients, and macromolecules (with Mw up to 1000 kDa) can freely pass through the hydrogel shell to feed cells inside the tube. Third, the hydrogel tubes confine the radial diameter of cell masses within the diffusion limit (∼400 μm) to ensure efficient mass transport during the whole culture period. Our previous study shows that the diffusion limit in 3D human cell culture is typically less than 500 µm^20,21,30–32,22–29^. Fourth, the hydrogel tubes also isolate hydrodynamic stresses in the culture vessels from cells. Lastly, the tubes provide free and uniform 3D microspaces that allow cells to interact with each other and proliferate (**Figure 1D**). Our previous studies showed that the free space is critical to achieving a high cell growth rate and yield^20,21,30–32,22–29^.

### C2C12 expansion in AlgTubes

We cultured C2C12 in AlgTubes (**Figure 2**). Day 0 image showed our seeding density was very low (1×10^6^ cells/mL hydrogel tube spaces). Only a few cells could be found in the amplified image on day 0. A low cell seeding density allows a large expansion fold per passage and is needed for industrial cell production. After 24 hours, all cells attached to the hydrogel tube’s inner surface with no or minor cell death. On day 4, cells formed a confluent monolayer. On day 7, multilayer cell masses were seen (green arrows). The dark-field image on day 7 showed white cell mass (orange arrows), indicating 3D multilayer cell masses. After 14 days, the dark-field images showed extensive 3D white cell masses (orange arrows), and some myotubes could be found in phase images (blue arrows). We found some locations of the tubes were bent (**Figure 2A**, red arrows) after 14 days, indicating the cell contraction force was bigger than the hydrogel elasticity. AlgTubes with stronger Young’s modulus can overcome this problem to further enhance the cell culture outcome in the future.

**Figure 2.**
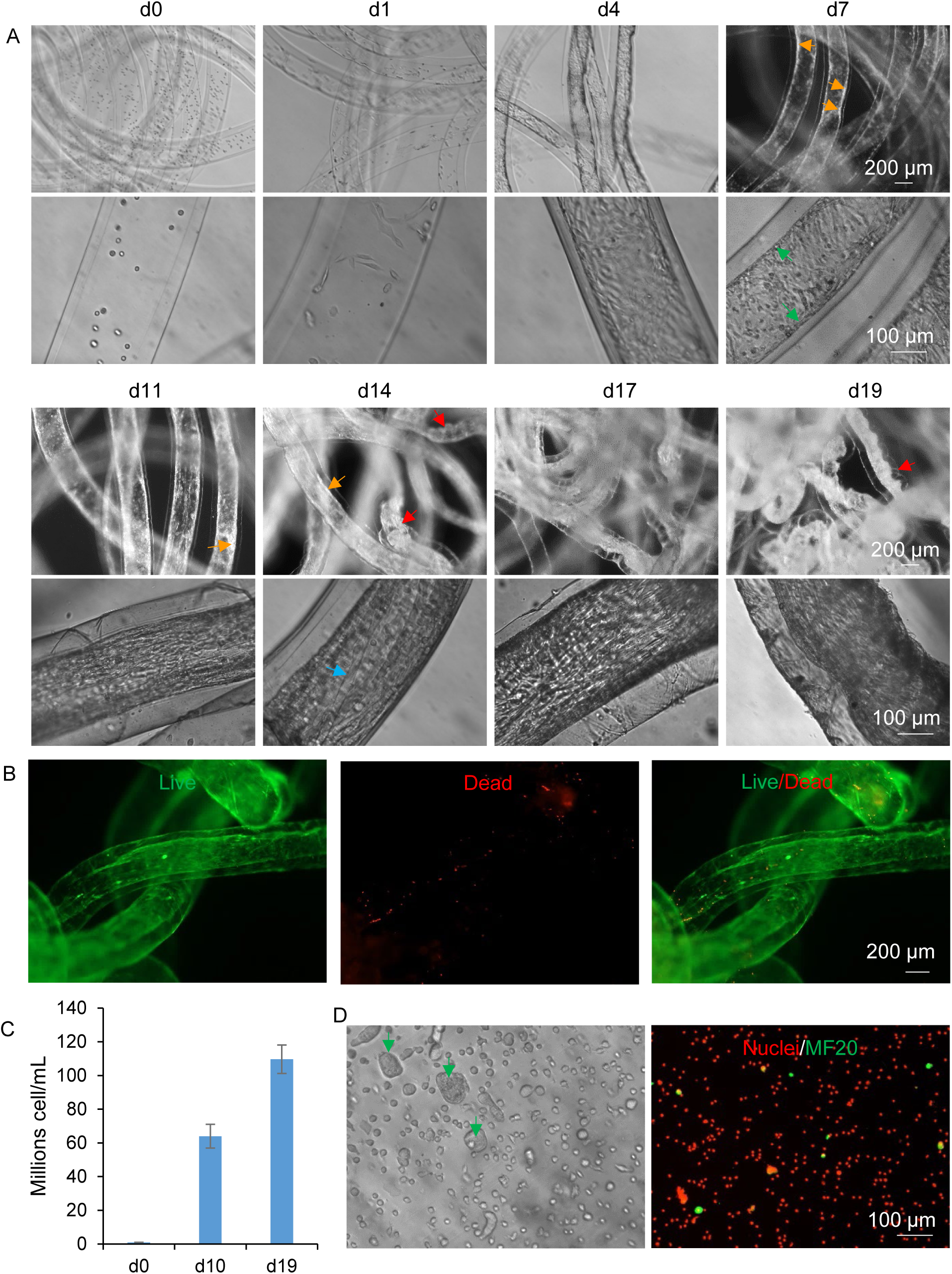
C2C12 expansion in AlgTubes. (A) Phase or dark-field pictures of C2C12 cells in AlgTubes on different days. Green arrows: multilayer cell masses; orange arrows: 3D cells mass; blue arrows: myotubes; red arrows: bent tubes. (B) Live/Dead cell staining of C2C12 in AlgTubes on day 19. (C) Cell densities on day 0, day 10 and day 19. (D) C2C12 cells released from

Live/Dead cell staining on day 19 showed that most were live cells (**Figure 2B**). The cell quantification shows that C2C12 cells reached ∼6.4×10^7^ and ∼1.1×10^8^ cells/mL hydrogel tube space on day 10 and day 19, respectively (**Figure 2C**). The released day 19 cells had healthy morphology. There were some large cell aggregates (**Figure 2D**, green arrows), which might be the contracted myotubes. Immunostaining on myofiber (MF20) confirmed the existence of a few myotubes (**Figure 2D**). Our results agree with the findings of the literature that C1C12 cells spontaneously differentiate to form myotubes at high density, even in the expansion medium.

### C2C12 differentiation in AlgTubes

Next, we studied if C2C12 cells could be differentiated into myotubes. Cells were expanded for 7 days in AlgTubes before initiating differentiation (**Figure 3**). Myotubes significantly increased after six days of differentiation (**Figure 3A**, blue arrows). Most cells were live based on the Live/Dead cell staining assay (**Figure 3B**). After differentiation for 12 days, the volumetric yield reached ∼1.2×10^8^/mL. We used immunostaining to evaluate the formed myotubes. The staining confirmed the 3D cell masses in the hydrogel tubes. There was still some space at the tube core (**Figure 3C and 3D**). A large percentage of cells, but not all, were MF20 positive (**Figure 3C**). The myotubes were large and aligned along the hydrogel tubes. Cells also expressed MyoD and PAX7, confirming the myo-cell lineage (**Figure 3D**).

**Figure 3.**
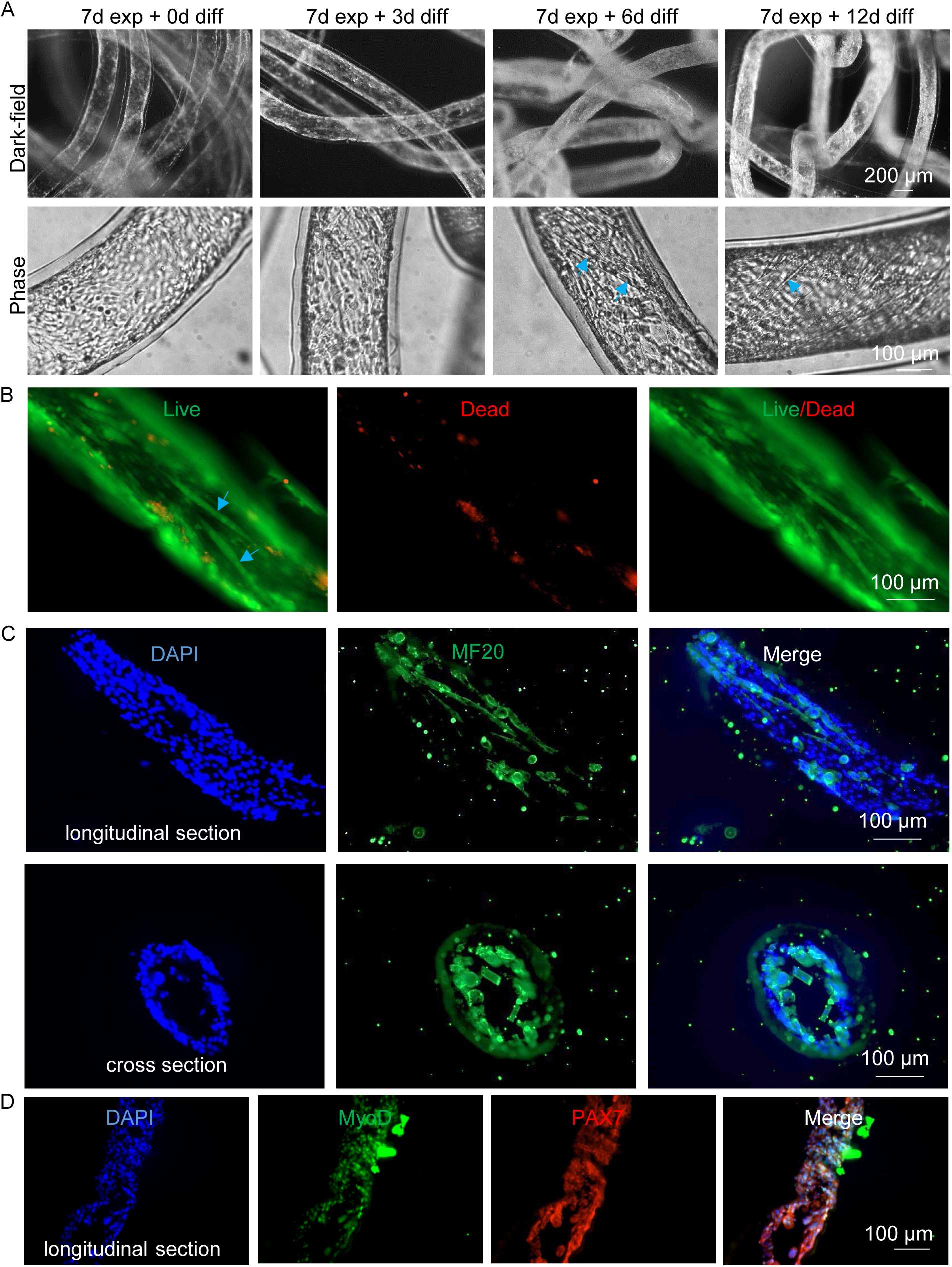
C2C12 differentiation in AlgTubes. C2C12 were cultured for 7 days in AlgTubes before differentiation for 12 day. (A) C2C12 cells in AlgTubes on different days. (B) Live/Dead cell staining on day 19. Blue arrows: Myotubes. (C) Immunostaining on 12 days. Cell fibers were fixed, cryosection and stained for myofibers (MF20) and DPAI (C) or stained for MyoD and PAX7 (D).

### C2C12 and D1 co-culture

Since AlgTubes were not fully populated by C2C12 cells (**Figure 3C**), we explored methods to enhance cell yield. Our previous research demonstrated that mesenchymal stem cells (MSCs) or fibroblasts significantly improve cell viability and yield in high-density cultures. Based on this, we conducted a co-culture experiment to assess whether mouse D1 MSCs could enhance the growth rate and yield of C2C12 cells in AlgTubes (**Figure 4**). D1 cells were pre-stained with the fluorescent cell-labeling dye DIO (green). C2C12 cells were seeded at 1 × 10⁶ cells/mL, along with 1 × 10⁵ D1 cells/mL in the hydrogel tubes. After 24 hours, cells attached to the hydrogel tubes and spread. By day 4, a confluent monolayer had formed on the inner surface. By day 7, significant cell growth resulted in the formation of multilayered cell masses (**Figure 4A**, green arrows). The presence of white cell masses in the dark-field image (Figure 4A, orange arrows) confirmed the formation of 3D multilayered structures. After 14 days of culture, extensive 3D cell masses were observed in the hydrogel tubes (orange arrows), along with some myotubes (blue arrows). The alginate hydrogel tubes bent under strong contractile forces exerted by the myotubes (red arrows).

**Figure 4.**
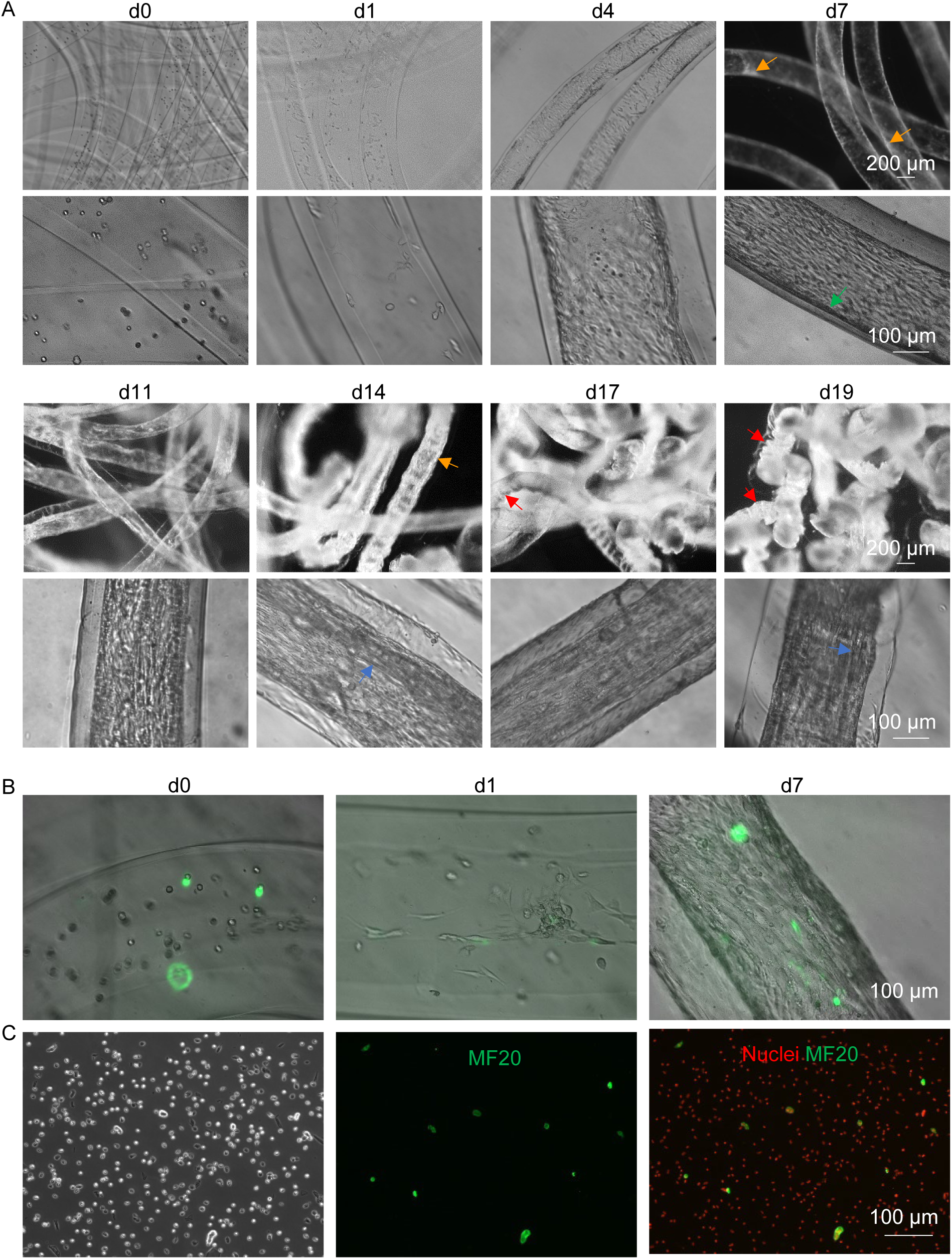
C2C12 and D1 cells co-culture in AlgTubes. (A) Cells in AlgTubes on different days. Green arrows: multilayer cell masses; orange arrows: 3D white cells mass; blue arrows: myotubes; red arrows: bent tubes. (B) Phase and fluorescent images of cells in AlgTubes. Green: D1 cells. (C) Cells were released from AlgTubes on day 19 and stained for MF20 and Nuclei.

To prevent D1 cells from dominating the culture, they were limited to 10% of the total cell population. Fluorescent imaging from day 0 to day 7 showed that D1 cells did not exhibit a growth advantage over C2C12 cells (**Figure 4B**). Cells maintained healthy morphologies when harvested and digested into single cells (**Figure 4C**). Immunostaining for myofibers (MF20) confirmed the presence of small numbers of myotubes, indicating that D1 cells did not inhibit spontaneous C2C12 differentiation.

We further investigated whether C2C12 cells could undergo differentiation in the presence of D1 cells (**Figure 5**). After 12 days of differentiation, immunostaining revealed that most cells were MF20-positive, confirming that D1 cells did not inhibit C2C12 differentiation (**Figure 5B**).

**Figure 5.**
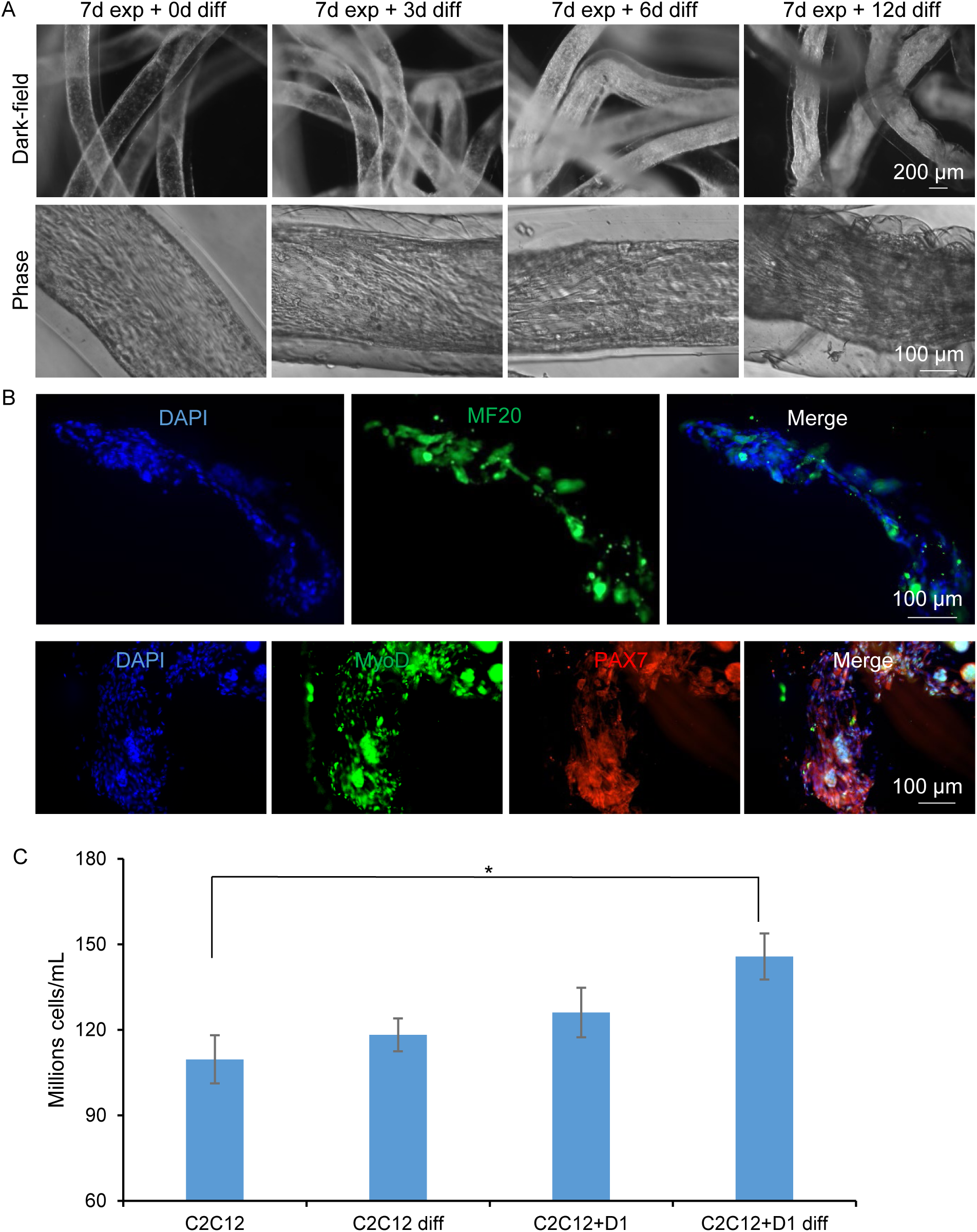
C2C12 and D1 co-differentiation in AlgTubes. C2C12 and D1 cells were cultured for 7 days in AlgTubes before differentiation. (A) Phase or dark-field pictures of cells on different days. (B) Day 19 cell mass cryosections were immunostained for MF20, MyoD and PAX7. (C) Quantifications under different conditions in AlgTubes on day 19.

While imaging suggested that D1 cells did not significantly alter the culture, cell counting showed that co-culture increased volumetric yield, particularly under differentiation conditions (**Figure 5C**). In the expansion medium, C2C12 cells reached approximately 1.1 × 10⁸ cells/mL of hydrogel tube space on day 19. Co-culturing with D1 cells increased the yield to ∼1.3 × 10⁸ cells/mL. In the differentiation medium, C2C12 cells alone reached ∼1.2 × 10⁸ cells/mL on day 19, while co-culturing with D1 cells boosted the yield to ∼1.6 × 10⁸ cells/mL. Therefore, adding D1 could improve the culture density, but not substantially. It should be noted that cell counting was performed using the Countess II cell counter. We believe the actual yields were higher than the reported values since the cell counter excluded large cell aggregates, which were frequently observed on day 19.

### QM7 expansion in AlgTubes

To confirm the broad applicability of RGD-AlgTubes for culturing animal cells, we cultured myoblasts from a second species, quail (QM7) (**Figure 6**). After 24 hours, most cells had attached to the hydrogel tube walls and exhibited a fibroblast-like morphology. By day 3, cells had expanded significantly, though a confluent monolayer had not yet formed. By day 6, multilayered cell masses were observed (**Figure 6A**, green arrows); however, some areas of the hydrogel’s inner surface remained uncovered (**Figure 6A**, blue arrow). This behavior differed from C2C12 cells (**Figure 2A**), which first formed a confluent monolayer before developing 3D cell masses. These findings suggest that quail myoblasts may have stronger cell-to-cell interactions but weaker adhesion to the hydrogel matrix than C2C12 cells. By day 9, dark-field images revealed extensive 3D white cell masses (**Figure 6A**, yellow arrows). Dense cell aggregates became more prominent by day 12 and were even more pronounced by days 15 and 18 (Figure 6A, red arrows). Quantification showed that QM7 cell yields on days 11 and 18 were approximately 1.2 × 10⁸ cells/mL (**Figure 6B**).

**Figure 6.**
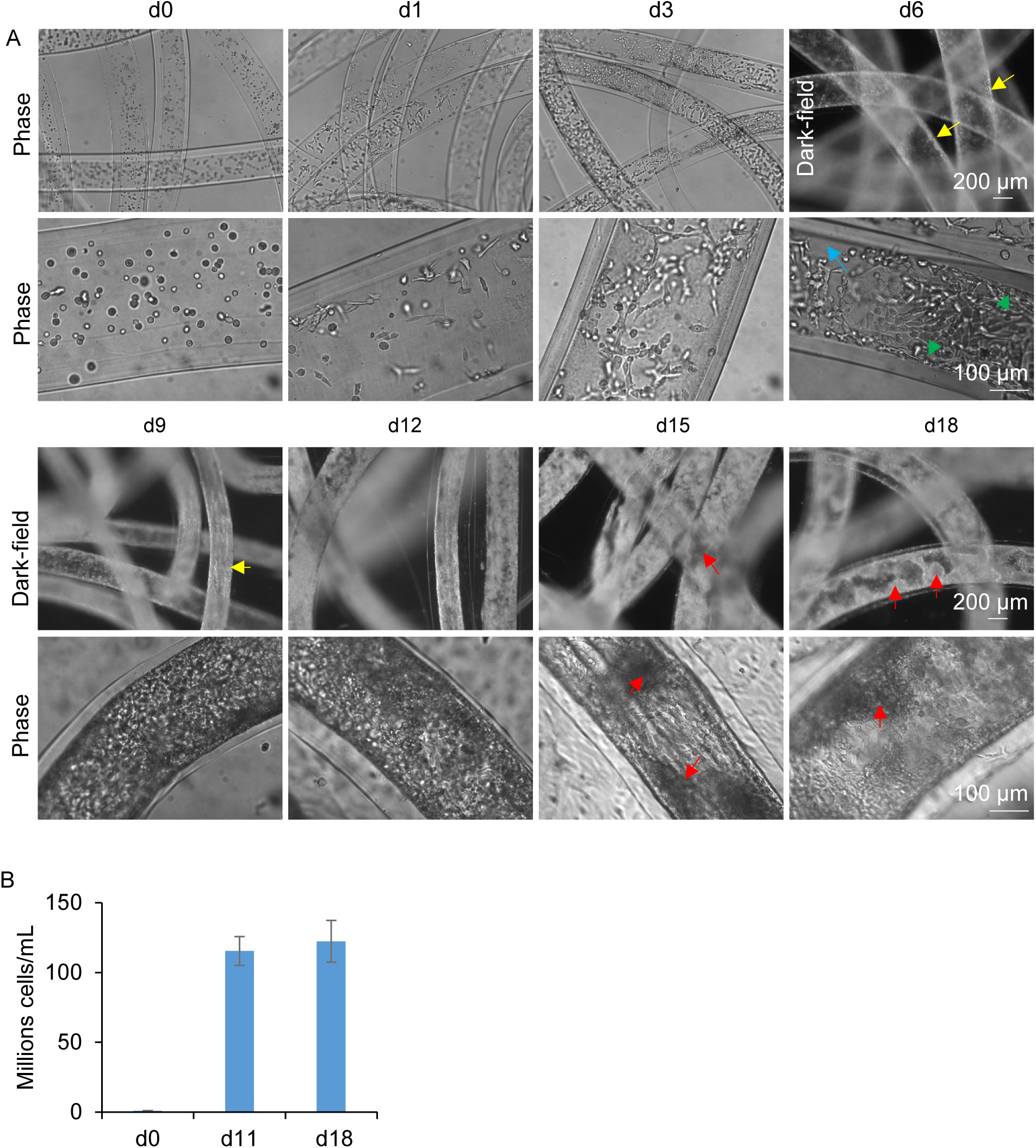
QM7 cells expansion in AlgTubes. (A) Phase or dark-field pictures of QM7 cells in AlgTubes on different days. Blue arrows: empty space; Green arrows: multilayer cell masses; yellow arrows: 3D white cells mass; red arrows: dense aggregates. (B) Cell yields on day 11 and 18.

Live/Dead staining on day 11 indicated that most released cells were viable, with only a few dead cells observed (**Figure 7A**). Staining of the whole-cell fiber revealed that dead cells were primarily located within dense multilayered regions. By day 18, Live/Dead staining showed increased cell death, particularly within large cell aggregates (**Figure 7B**). Despite continued cell proliferation between days 11 and 18, the overall yield remained similar due to the rise in cell death (**Figure 6B**). Compared to C2C12 cells, QM7 cultures exhibited higher levels of cell death by day 18, likely due to their weaker adhesion to the hydrogel and tendency to form aggregates.

**Figure 7.**
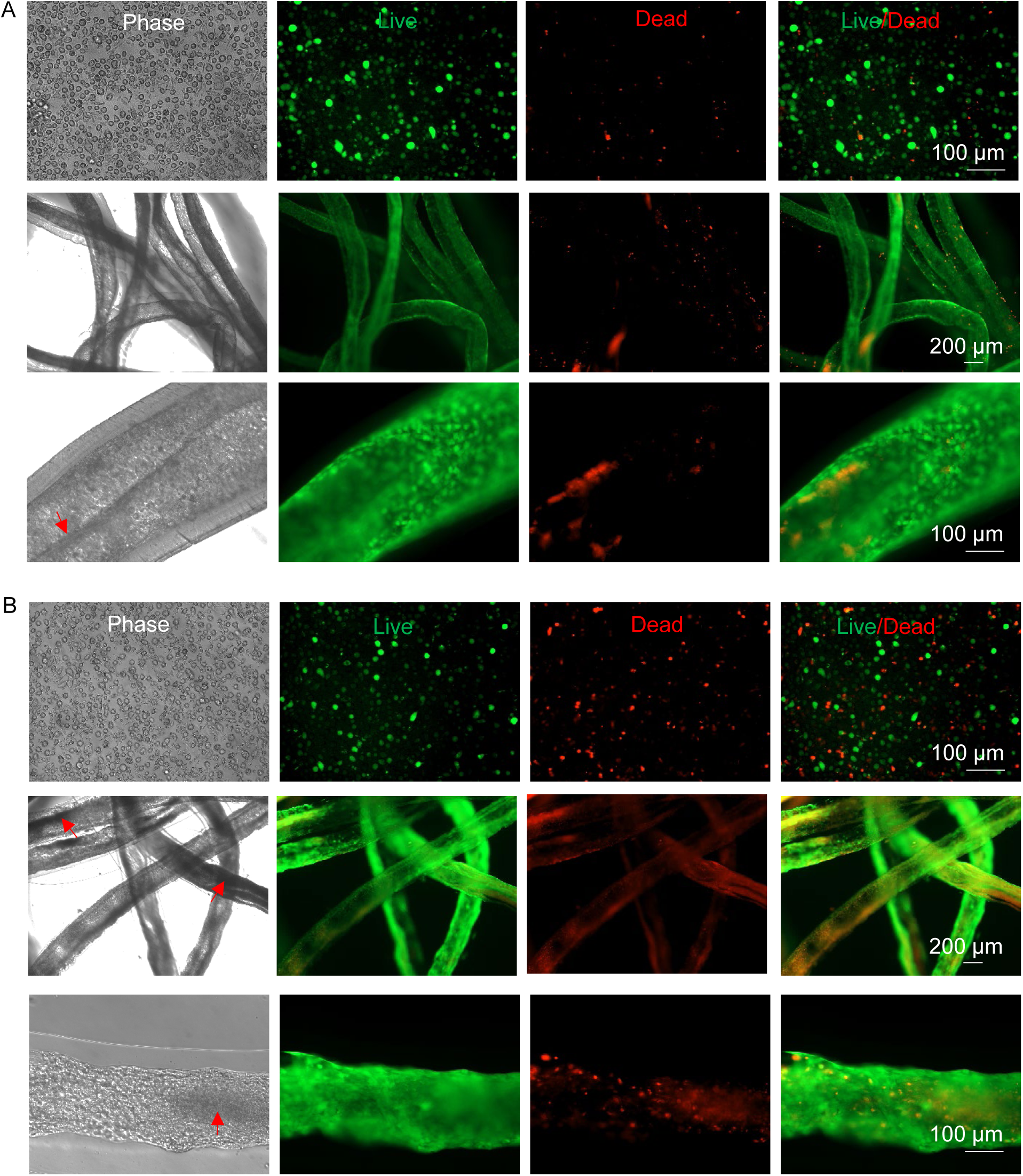
Live/Dead cell staining of QM7 cultured in AlgTubes on day 11 (A) and day 18 (B). The first row of (A) and (B): released and digested cells on day 11 and day 18. Red arrows: dense aggregates.

Our results demonstrate that C2C12 and QM7 cells behave differently in AlgTubes. While C2C12 cells formed a confluent monolayer before developing 3D structures, quail cells preferentially formed 3D aggregates. To determine whether this difference was due to the AlgTubes or an intrinsic characteristic of the cells, we cultured and differentiated C2C12 and QM7 cells in traditional 2D culture dishes (**Figures S1 and S2**). By day 5, both cell types had formed myotubes. By day 11, C2C12 cells had generated more myotubes, whereas quail myotubes exhibited an aggregative morphology. These findings confirm that AlgTubes did not alter the intrinsic properties of the cells.

### Quail cell differentiation in AlgTubes

We further examined QM7 cell differentiation in AlgTubes. Quail cells were first expanded for six days, reaching 80–90% confluence before initiating differentiation (**Figure 8**). During differentiation, cells continued to proliferate and formed 3D cell masses. Myotubes became visible after three days of differentiation (**Figure 8A**). Live/Dead staining revealed that dead cells were primarily located within large, dense aggregates (**Figure 8B**, red arrows). Immunostaining confirmed the presence of myotubes aligned along the hydrogel tubes. Additionally, the myotubes expressed MyoD and PAX7 (**Figure 8C**).

**Figure 8.**
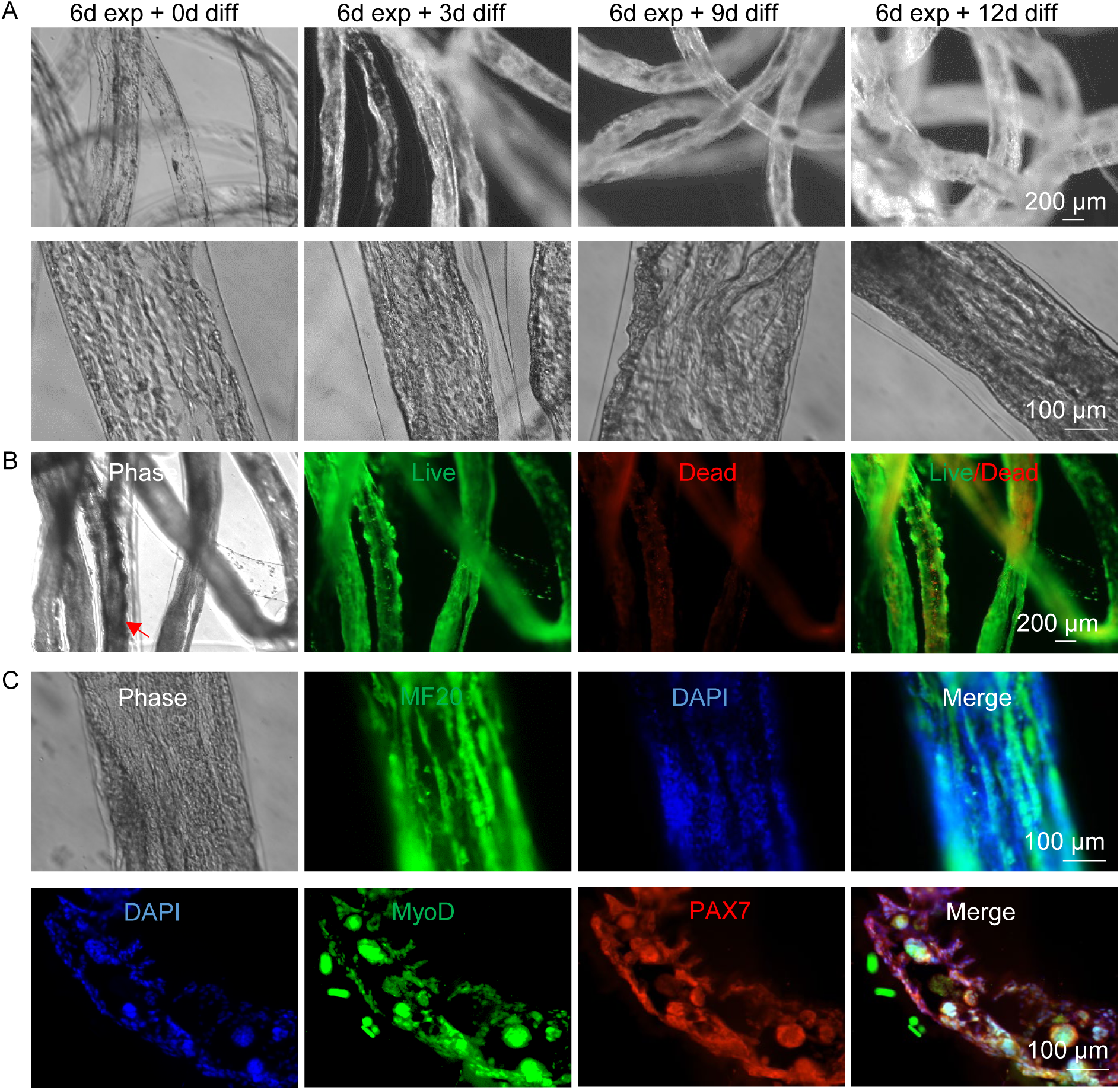
QM7 differentiation in AlgTubes. QM7 were first expanded for 6 days in AlgTubes before differentiation. (A) Phase or dark-field pictures of QM7 cells on different days. Day 18 cells were harvested for live/dead staining (B) and fixed for MF20, MyoD and PAX7 immunostaining (C).

### QM7 and 3T3 cell co-culture

Like C2C12, we performed a co-culture of QM7 and D1 cells, yielding comparable results (**Figures S3 and S4**). Fibroblasts, another type of stromal cell, are known to enhance the survival of other cell types. Therefore, we conducted co-culture experiments to evaluate whether mouse 3T3 fibroblasts could improve the growth rate and yield of QM7 cells in AlgTubes (**Figure 9**). After 24 hours, cells attached to the hydrogel tubes with minimal cell death. Dense aggregates developed by day 12 and continued to expand. By day 18, Live/Dead staining indicated dead cells primarily localized within large, dense aggregates (**Figure 9B**).

**Figure 9.**
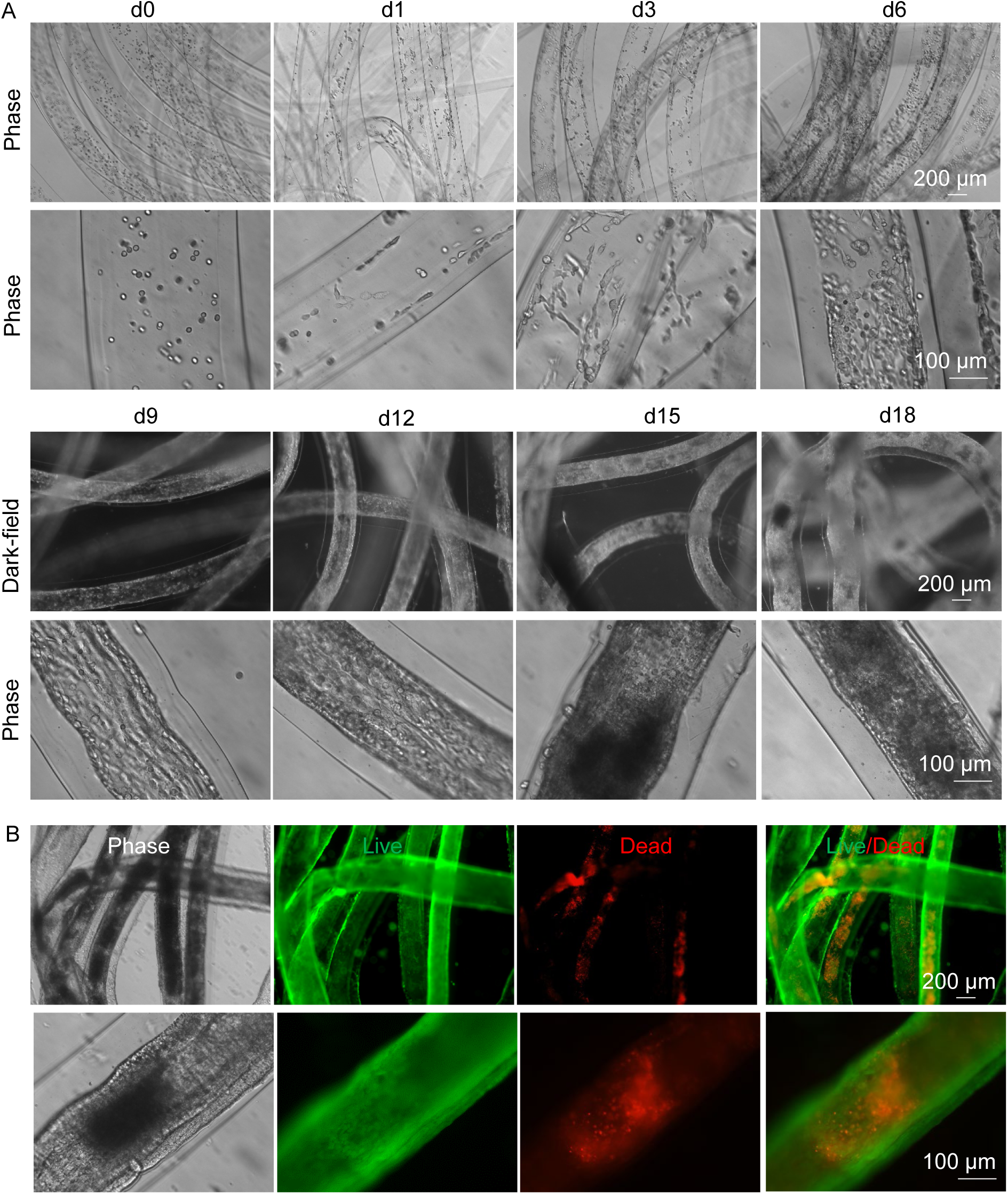
QM7 and 3T3 co-culture in AlgTubes. (A) Phase or dark-field pictures of cells in AlgTubes on different days. (B) Live dead cell staining on day 18.

Differentiation was initiated after six days of QM7 and 3T3 co-culture in AlgTubes. Myotubes appeared after six days of differentiation (**Figure 10A**), accompanied by the formation of dense aggregates. By day 18, Live/Dead staining still showed substantial cell death within these aggregates (**Figure 10B**). Immunostaining confirmed that many cells were MF20-positive, verifying myotube formation. We quantified the cell yield under different conditions (**Figure 10C**). Overall, QM7 cells in AlgTubes achieved a final yield of over 1.1 × 10⁸ cells/mL.

**Figure 10.**
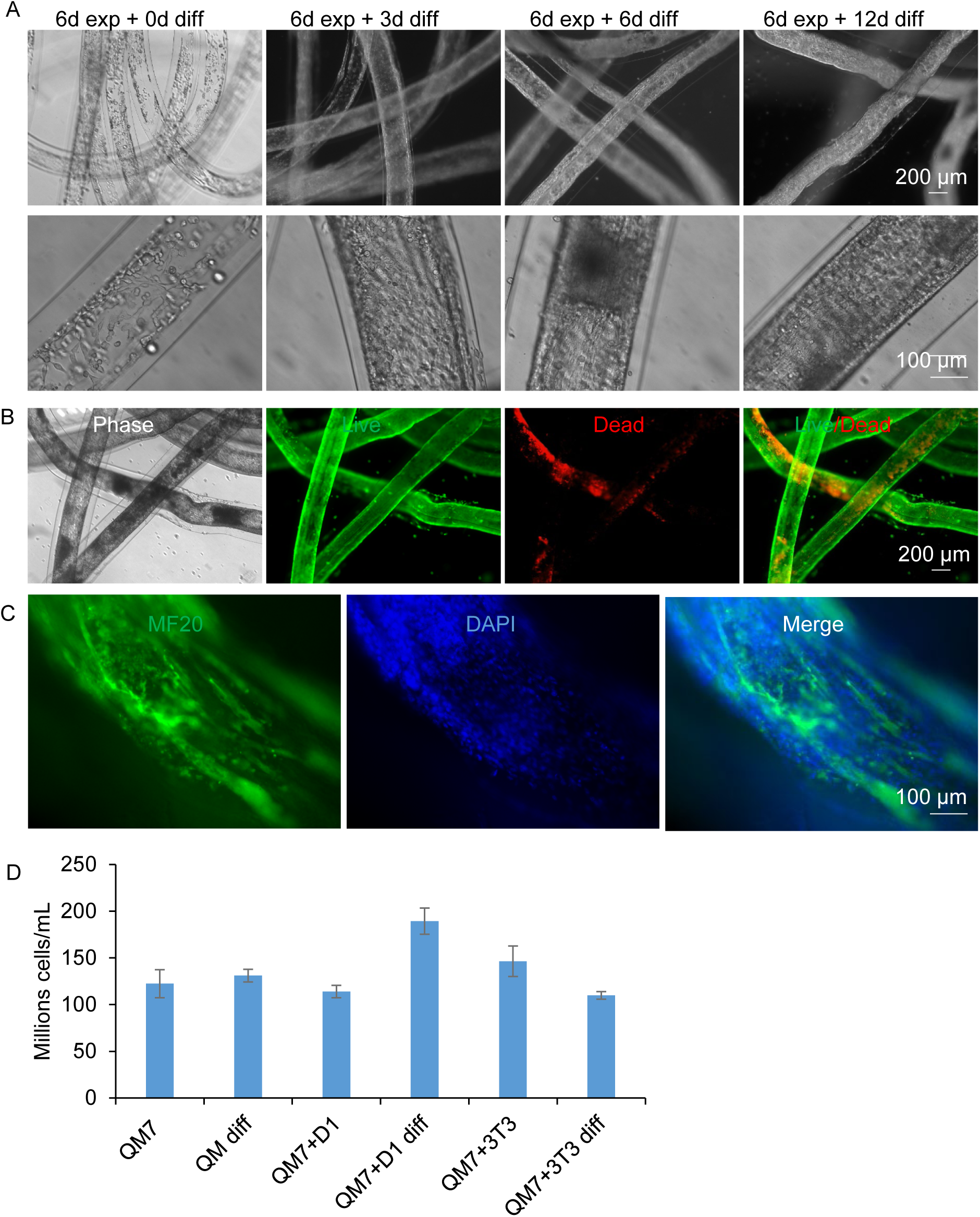
QM7 and 3T3 cells differentiation in AlgTubes. QM7 and D1 cells were first expanded for 6 days in AlgTubes before differentiation. (A) Phase or dark-field pictures of cells in AlgTubes on different days. Day 18 cells were harvested for live/dead staining (B) and fixed for MF20 immunostaining (C). (D) Quantifications under different conditions on day 19

### X9 expansion in AlgTubes

Adipocytes are a crucial component of meat composition. To assess whether our system could support the production of pre-adipocytes or adipocytes, we cultured mouse pre-adipocyte X9 cells (**Figure 11**). X9 cells were seeded at a low density (1 × 10⁶ cells/mL). After 24 hours, most cells were attached to the hydrogel tube walls, with minimal cell death. Unlike C2C12 and QM7 cells, X9 cells expanded very slowly. Cell numbers on day 3 remained similar to day 1, and a confluent monolayer had not yet formed even after 10 days. By day 13, most areas had reached confluence, though only a few multilayered cell masses were observed (**Figure 11A**, green arrows). Even after 19 days of culture, most of the hydrogel tubes’ inner surfaces remained covered by a single layer of X9 cells, with no large cell masses detected. The final yield of X9 cells was ∼3.5 × 10⁷ cells/mL on day 19 (**Figure 11B**). Live/Dead staining confirmed that most cells remained viable (**Figure 11C**).

**Figure 11.**
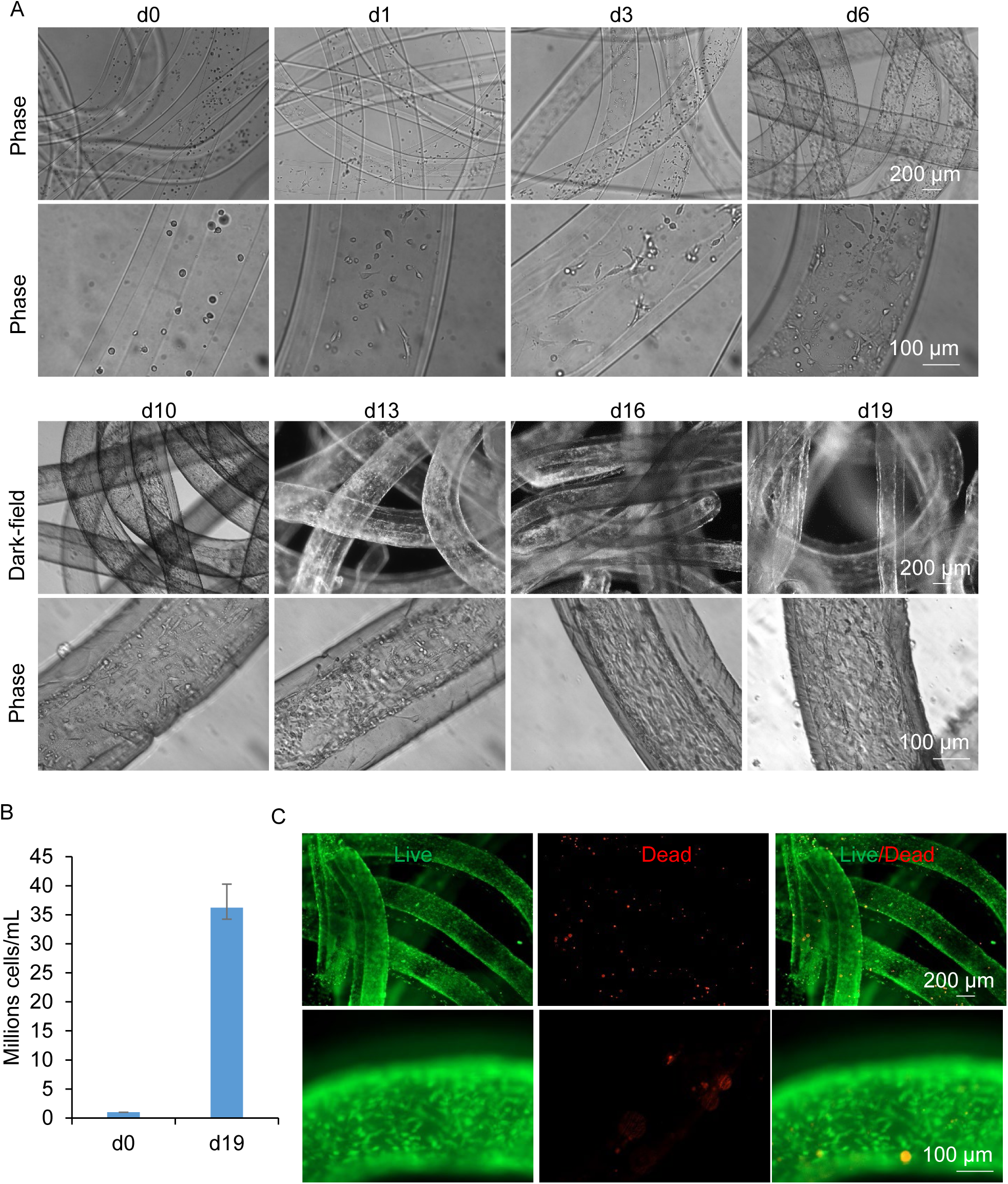
X9 cells expansion in AlgTubes. (A) Phase or dark-field pictures of X9 cells on different days. (B) Cell quantification on day 0 and day 19. (C) Live/Dead cell staining on day 19.

## Discussion

Cultured meat holds substantial promise for addressing the growing global demand for food in a sustainable and ethical manner, given the environmental and ethical issues associated with traditional livestock farming. However, a key challenge facing cultured meat production is achieving industrial-scale cell cultivation efficiently and economically. Although three-dimensional (3D) suspension culturing methods, such as stirred-tank bioreactors (STRs), have revolutionized the production of therapeutic proteins, their effectiveness in scaling up stem cells and cultured meat production is limited by several critical issues^9,11,34–36^.

First, mammalian cells exhibit strong cell-to-cell adhesion, leading to the formation of large aggregates^37–41^. These aggregates frequently surpass the diffusion limit of nutrients and oxygen (approximately 400 µm), causing insufficient nutrient delivery, reduced cell proliferation, apoptosis, and unintended differentiation^41,42^. Second, the agitation employed in STRs generates complex and variable hydrodynamic conditions, which induce harmful shear stresses, particularly near bioreactor walls, further reducing cell viability and overall culture efficiency^39–41^. For example, previous research, including our own, indicates that human pluripotent stem cells (hPSCs) typically exhibit less than a 10-fold expansion per passage in suspension culture, achieving yields of fewer than 3×10⁶ cells/mL^9,11,34–36^. These cells occupy less than 0.5% of the culture volume, highlighting the inefficiencies of current methods. Additionally, hydrodynamic conditions within STR systems are sensitive to multiple variables, including bioreactor geometry, impeller design, medium viscosity, and agitation speed^41,43^. Variations in hydrodynamic conditions significantly impact cell culture outcomes, resulting in inconsistent performance between batches and difficulties scaling from laboratory to industrial volumes ^39–41,43–45^.

To address these limitations, we modified our previously established AlgTubes technology by incorporating RGD peptides, enabling the efficient cultivation of anchorage-dependent cells, including myoblasts and adipocytes. Our results demonstrate that RGD-modified AlgTubes significantly enhance cell density or yield compared to conventional STR-based methods. Differences in cell behavior within AlgTubes provided insights into optimizing hydrogel conditions for diverse cell types. While C2C12 cells formed confluent monolayers before forming multilayered masses, QM7 cells tended to aggregate more strongly, resulting in cell death within dense masses after extended culture periods. Adjustments in RGD concentration or hydrogel stiffness could potentially reduce aggregation and improve outcomes for cells with strong intrinsic cell-to-cell interactions.

Co-culture strategies further improved outcomes. Co-culturing mouse MSCs (D1) or fibroblasts (3T3) with myoblasts moderately enhanced cell viability and yield, particularly during differentiation conditions. Importantly, co-culture did not negatively affect differentiation capacity, as evidenced by the presence of robust myotube formation. These results suggest that AlgTubes is compatible with complex co-culture systems, which may be beneficial for recreating the heterogeneity and structure of natural meat.

Our system was less effective for adipocytes (X9 cells), which proliferated more slowly, resulting in yields of around 3.5×10⁷ cells/mL after 19 days. While viability was maintained, optimization such as adjusting nutrient formulations or incorporating supportive stromal cells might further enhance adipocyte growth and achieve yields suitable for cultured meat production.

## Conclusion

Our findings demonstrate that RGD-modified AlgTubes represent a highly effective platform for the scalable culture of anchor-dependent animal cells essential for meat production. AlgTubes substantially outperform current state-of-the-art cell culture methods in terms of cell yield. By achieving cell densities exceeding 100 million cells per milliliter, AlgTubes substantially reduces culture volume, labor requirements, reagent costs, equipment needs, facility space, and overall manufacturing expenses, making large-scale production feasible and practical.

Future studies should optimize AlgTube parameters—such as RGD peptide concentration, hydrogel stiffness, and medium composition—to further enhance performance across different cell types, particularly adipocytes. Additionally, scale-up validations are necessary to confirm commercial viability and sustainability. Ultimately, the AlgTube platform holds great promise for revolutionizing meat production, offering an environmentally friendly and ethically sound solution to global food security challenges.

## Author Contributions

Y.L. conceived the idea and supervised the research. Y.L., S.W., O.W., and L.H. designed the experiments. O.W. and L.H. conducted experiments. Y.L., S.W., O.W., L.H. C.D., L.X. A.W. carried out data analysis and manuscript preparation.

## Data Availability

The authors confirm that the data supporting the findings of this study are available within the article.

---

**Figure S1.**
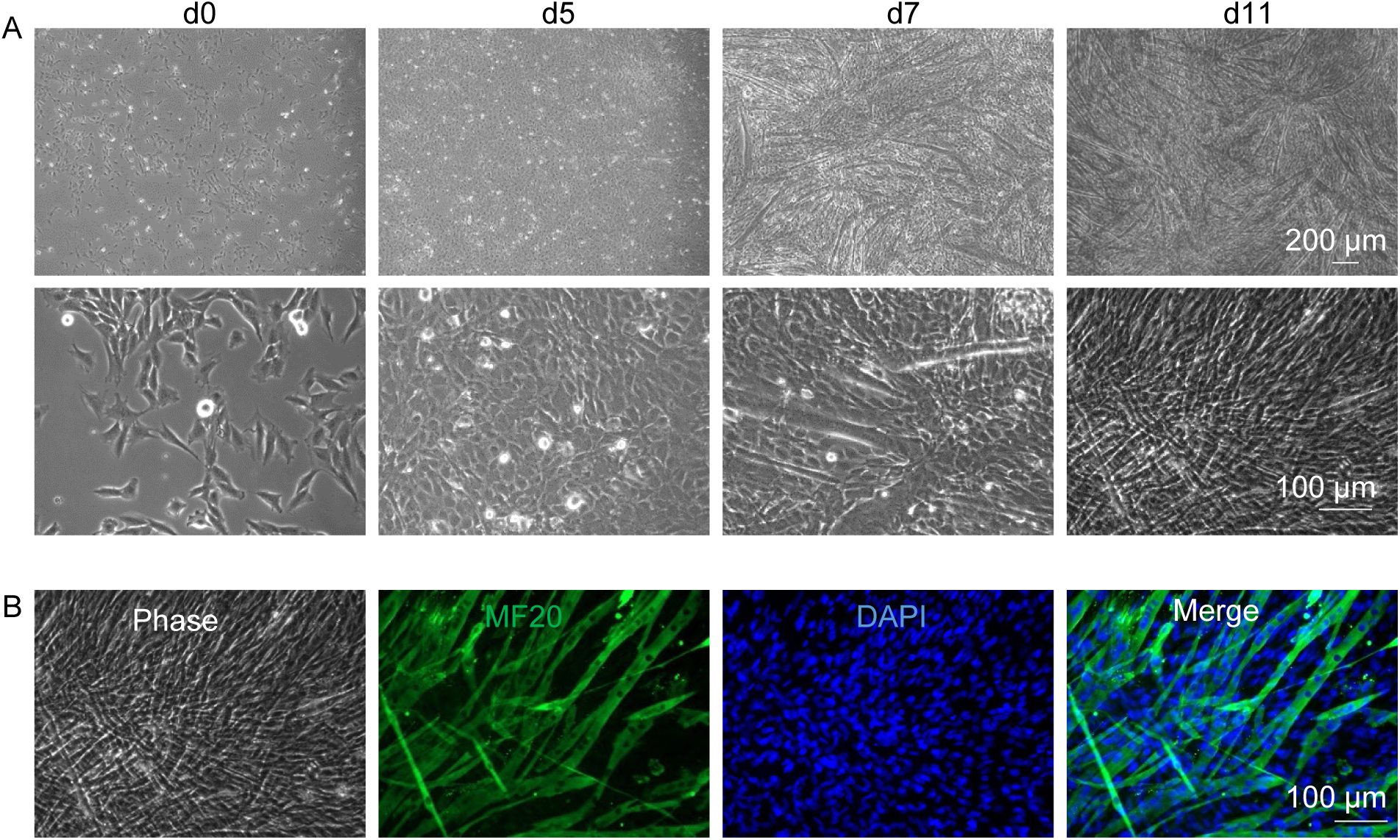
C2C12 differentiation in 2D culture. Differentiation started on day 3. (A) Phase images. (B) Immunostaning for MF20 on day 11.

**Figure S2.**
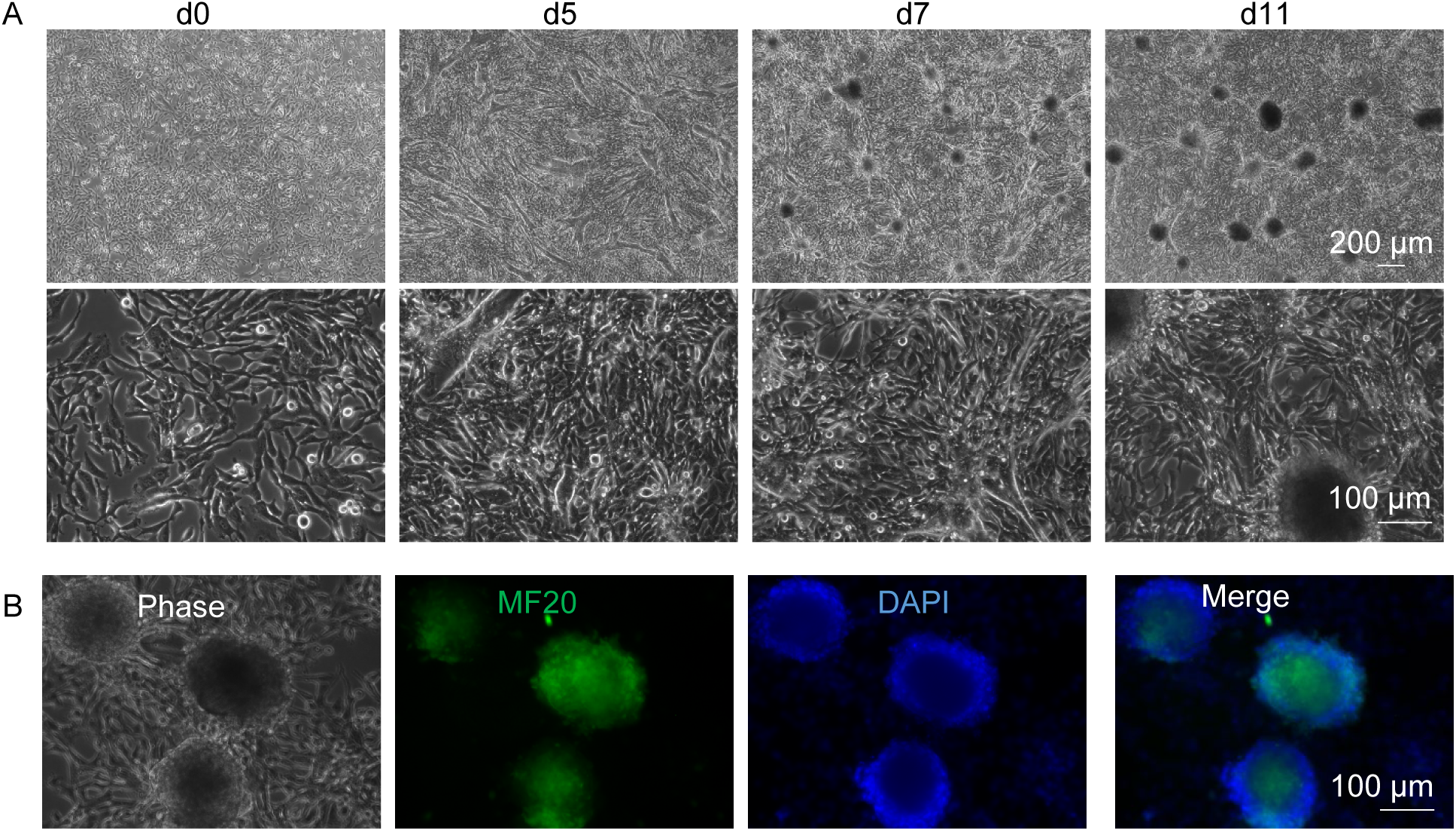
QM7 differentiation in 2D culture. Differentiation started on day 3. (A) Phase images. (B) Immunostaining for MF20 on day 11.

**Figure S3.**
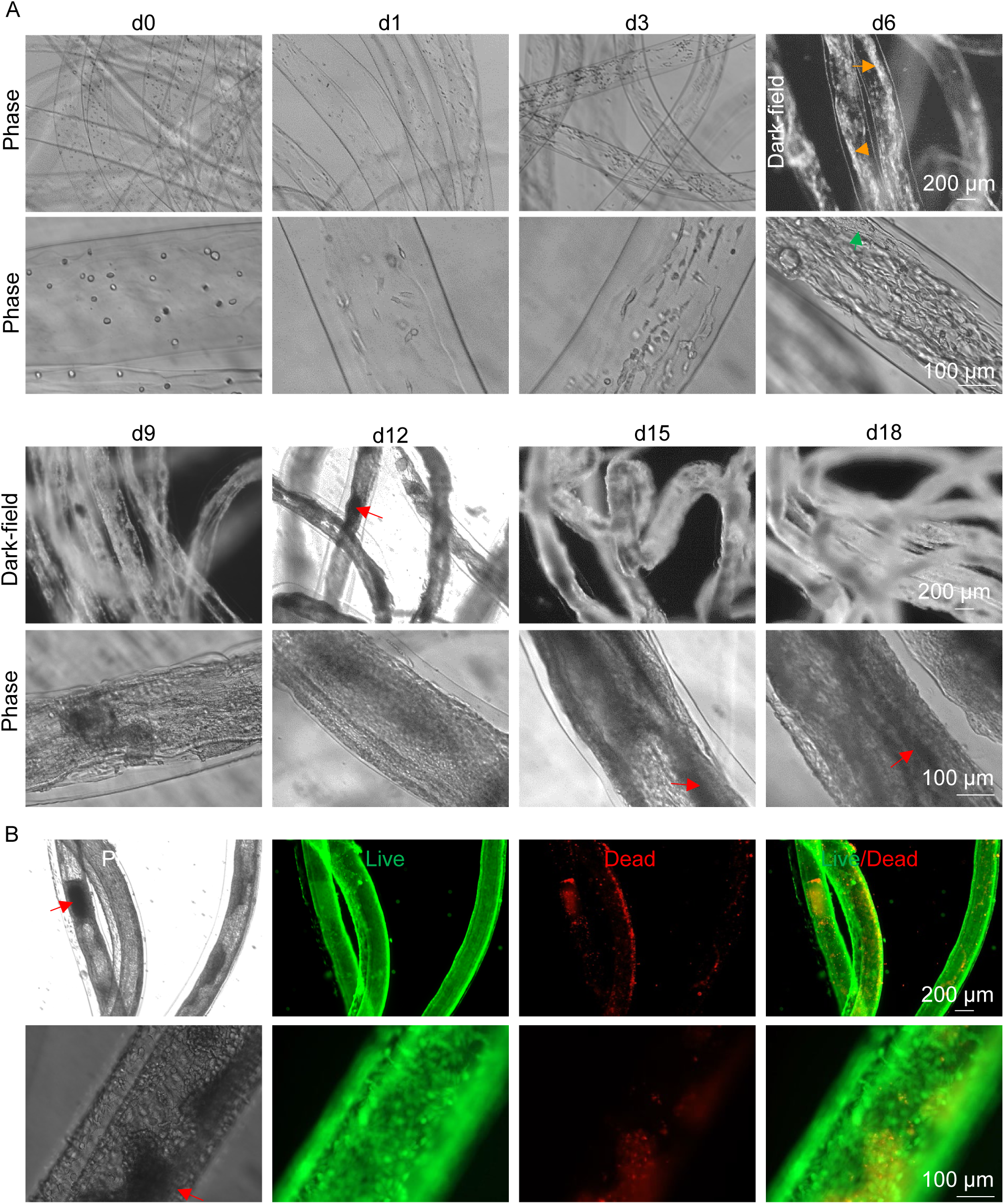
QM7 and D1 cells expansion in AlgTubes. (A) Phase or dark-field pictures of cells in AlgTubes on different days. (B) Live dead cell staining on day 18. Green arrows: multilayer cell masses; orange arrows: 3D white cells mass; red arrows: dense aggregates

**Figure S4.**
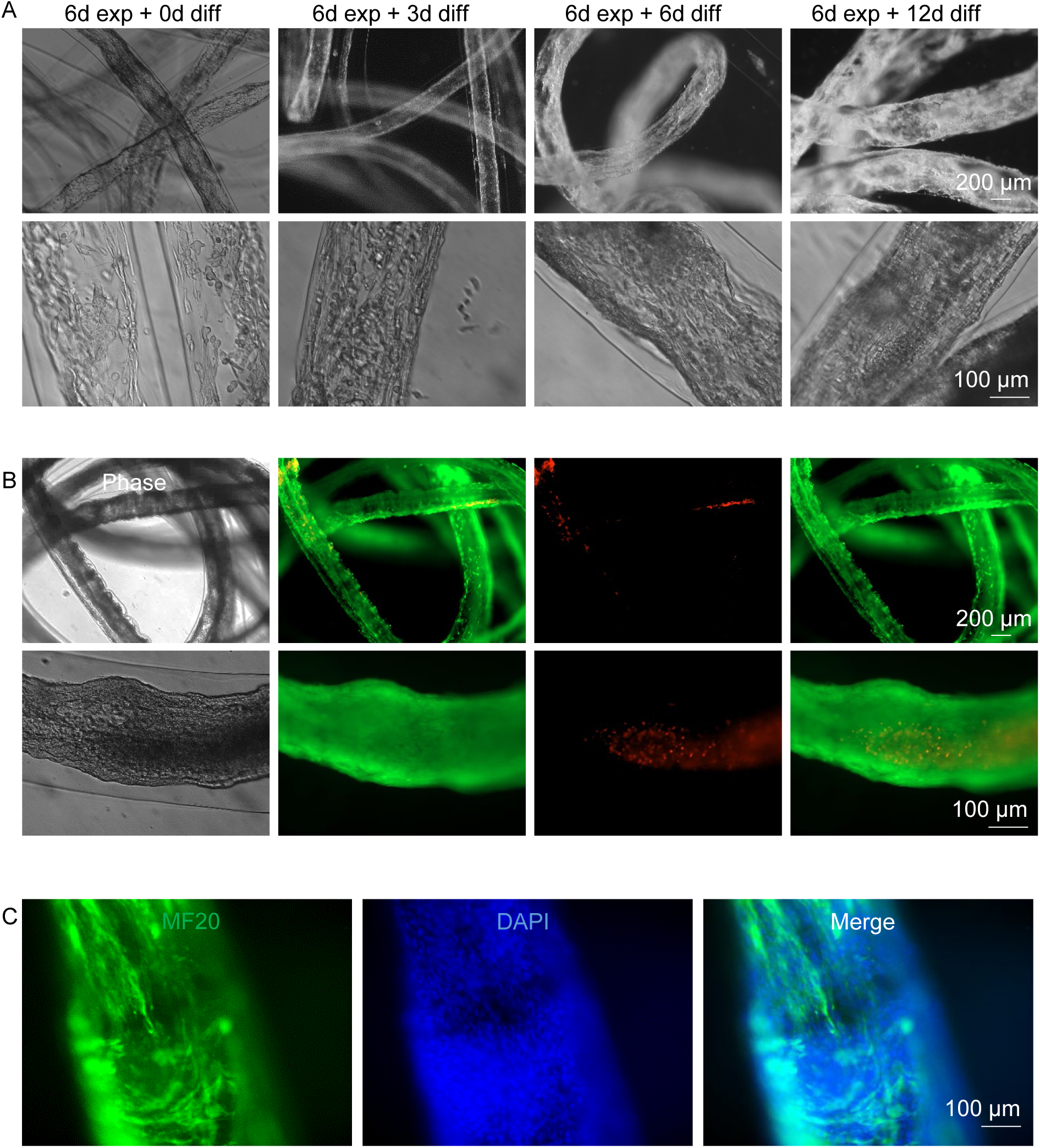
QM7 and D1 cell differentiation in AlgTubes. QM7 and D1 cells were first expanded for 6 days. (A) Phase or dark-field pictures of cells in AlgTubes on different days. Day 18 cells were harvested for live/dead staining (B) and fixed for MF20 immunostaining (C).

